# PREDICTING HISTORIC, CONTEMPORARY, AND FUTURE DISTRIBUTIONS OF *CULEX CORONATOR* FOLLOWING RAPID RANGE EXPANSION

**DOI:** 10.64898/2026.08.05.742963

**Authors:** Olivia R. Magaletta, Amely M. Bauer, Yoosook Lee, Panpim Thongspring, Lindsay P. Campbell

## Abstract

Invasive mosquito species pose substantial risks to human and animal health. Since 2004, *Culex coronator*, a mosquito vector species of public health concern, has shown rapid range expansion within the United States, spreading from a historically limited distribution in southern Texas to across the Gulf Coast region and into eastern and mid-Atlantic states. However, changes in environmental suitability associated with this expansion across historical, contemporary, and future climate conditions have not been evaluated. Here, we used species distribution models (SDMs) to compare predictions of abiotic suitability for *Cx. coronator* under historic (1960–1989) and recent (2000–2024) climate conditions calibrated on the historical range in the United States. We also created a contemporary SDM based on occurrence records prior to and following species range expansion (1960–2024), and further, to predict potential distributions under current and future climate conditions.

Models calibrated on the historical range predicted only modest changes in suitability along the Gulf Coast region and failed to identify large areas of the humid subtropical eastern United States that are now occupied. In contrast, the contemporary model predicted widespread suitability across much of the southern and eastern United States. Future projections under the mid-range SSP3 scenario predicted increasing suitability at higher latitudes and elevations. Across all models, suitability was consistently low in arid and semi-arid regions, including along the historical western range limit, suggesting that moisture availability may constrain *Cx. coronator* distributions. Together, these results highlight the need to incorporate updated occurrence records when modeling invasive mosquito species to strengthen surveillance and control strategies.

## Introduction

Invasive mosquito species are responsible for transmitting multiple pathogens affecting human and animal health [1]. The Gulf Coast region of the southern United States, including Texas, Louisiana, Mississippi, Alabama, and Florida, has seen persistent introduction or movement of invasive mosquito species, with several species moving eastward from Texas toward Florida, making this region a priority for ongoing surveillance and proactive vector management [2–7]. Among these species, *Culex coronator* has exhibited one of the most notable and sustained range expansions. *Culex coronator* is a multivoltine generalist species, occupying diverse habitats [8–10]. First documented in the United States in 1920 in San Benito, Texas [11], the species remained largely restricted to South and Central America, Mexico, and southern Texas throughout much of the 20th Century [12], with only transient detections elsewhere in the southwestern United States [12–21]. This pattern changed in 2004, when established populations were recorded in Louisiana and Mississippi [22,23]. Since then, *Cx. coronator* has expanded substantially across portions of the humid subtropical region of the eastern United States (U.S.), now occurring as far north as Virginia, south to Florida, and west to New Mexico, and is currently reported in at least 14 states [16,24–29]. Given the rapid and ongoing expansion, understanding environmental suitability across historical, contemporary, and future conditions is essential for anticipating the potential distribution of this species for improved vector control to reduce public health risk.

*Culex coronator* is a public health concern due to its demonstrated ability to vector multiple pathogens, and several pathogens have also been detected in wild field collections, including West Nile virus and St. Louis encephalitis virus [30–32]. In addition, *Cx. coronator* feeds on both mammalian and avian hosts [8,9,33], demonstrating potential to serve as a bridge vector between wildlife and human transmission cycles [30–36]. The species can be collected year-round in some areas and can overwinter as an adult, making it a candidate for further range expansion under ongoing environmental change [27].

The Gulf Coast region is classified as humid subtropical based on the Köppen-Geiger climate classification, which covers a large portion of the eastern United States from Texas to the Atlantic Coast and northward from Kansas to southern parts of New Jersey [37]. Global mean temperature has risen approximately 1.1°C above pre-industrial levels, with warming since 1970 occurring faster than in any other 50-year period [38]. Future global climate projections suggest temperature is likely to increase > 2°C relative to pre-industrial levels by the end of the century [38–40]. The Gulf Coast region is also likely to see more extreme weather events, including more severe storms and heavier rainfall [38–40]. Together, these factors may alter environmental suitability for some mosquito species, with the potential to facilitate further range expansion.

The geographic distributions of mosquito species are shaped by multiple interacting abiotic and biotic factors [41,42]. Physical barriers can limit dispersal, while temperature, humidity, and the availability of larval habitats provide key environmental constraints limiting potential range. Biotic interactions such as competition, predation, and resource availability further influence local persistence, whereas abiotic climatic conditions impose broader limits on potential range. As ectothermic insects, mosquitoes are highly sensitive to temperature, and their small body size makes them especially vulnerable to desiccation from low humidity [43]. Consequently, abiotic climate variables are widely used to model and predict potential distributions of mosquito species.

Species distribution models (SDMs) are a set of models that correlate the combination of environmental conditions where a species has been observed to predict where environments may be suitable across a broader geographic area or different time period [44–46]. These models have been used to predict potential distributions of multiple medically important, non-native, and invasive vectors across regional to global scales [47–51]. Species distribution models have also been used to predict changes in potential distributions of vector species under climate change scenarios, often demonstrating possible shifts to northern or southern latitudes away from the equator or to higher elevations [52–57]. Despite the recent and ongoing range expansion of *Cx. coronator*, SDMs have not been used to examine the potential distribution of this species during historic, current, or under projected future climate conditions.

As *Cx. coronator* continues to expand geographically, understanding change in environmental suitability and anticipating future range potential is critical for effective surveillance and control. The objectives of this study are to: (1) create a historical species distribution model (SDM) calibrated with occurrence data before the 2004 range expansion (1960 – 1989) and projected to recent climate conditions (2000 - 2024) to assess changes in predicted suitability along the Gulf Coast; (2) develop a contemporary SDM including post-expansion records (1960 – 2024); and (3) project potential distributions under future climate scenarios (2041–2060). We expect that environmental suitability has increased across the Gulf Coast under recent climate conditions, and that future projections will indicate continued expansion northward and into higher elevations.

## Materials and Methods

### Occurrence Data

Georeferenced occurrence data for *Cx. coronator* were assembled from the Global Biodiversity Information Facility (GBIF) and from the scientific literature. GBIF-mediated data were downloaded on 17 December 2024 (https://doi.org/10.15468/dl.wf953x) [58] and imported into OpenRefine v3.8-beta1 [59] to remove duplicate records. The cleaned GBIF records were then formatted, using ‘tidyverse’ (V 2.0.0) functions in R v4.3.2 [60,61] before combining them with the georeferenced occurrence data identified in scientific literature. Next, the combined occurrence data were mapped and spatially thinned to a minimum distance of 0.25 decimal degrees to reduce sampling bias that could lead to redundancy in environmental combinations in model calibration [62] using functions available in the ‘ntbox’ (V 0.7.2) package in R [63]. We then created two occurrence data sets in preparation for SDM calibration: First, a data set containing occurrence records from 1960 to 1989 to create a historical data set for model calibration prior to geographic expansion. Second, all occurrence records (1960 to 2024), prior to and after range expansion, were assembled for a contemporary model calibration.

### Calibration Region

A user-defined calibration region is a critical element of SDMs [64]. Several methods can be used to define this region, including the creation of a convex hull around occurrence records, as well as taking into consideration areas accessible to a species across a geographic area and time period [64,65], outlined in the Biotic-Abiotic-Movement (BAM) framework [66]. Here, we defined the calibration region (**‘M’**) including areas accessible to *Cx. coronator* across South and Central America. This region excluded areas west of the Andes Mountains in South America, which served as a geographic barrier, and southern portions of the continent where occurrence records were not observed. We also included Central America and Mexico, and then constrained the northern and eastern boundaries to portions of California, Arizona, New Mexico, and Texas based on historical distributions (1960 - 1989) (Figure 1A). For the updated contemporary model, we included all regions within the historic limit, but constrained the northern limit to include the northern occurrence records and the humid subtropical climate range in the U.S. (Figure 1B).

**Figure 1.**
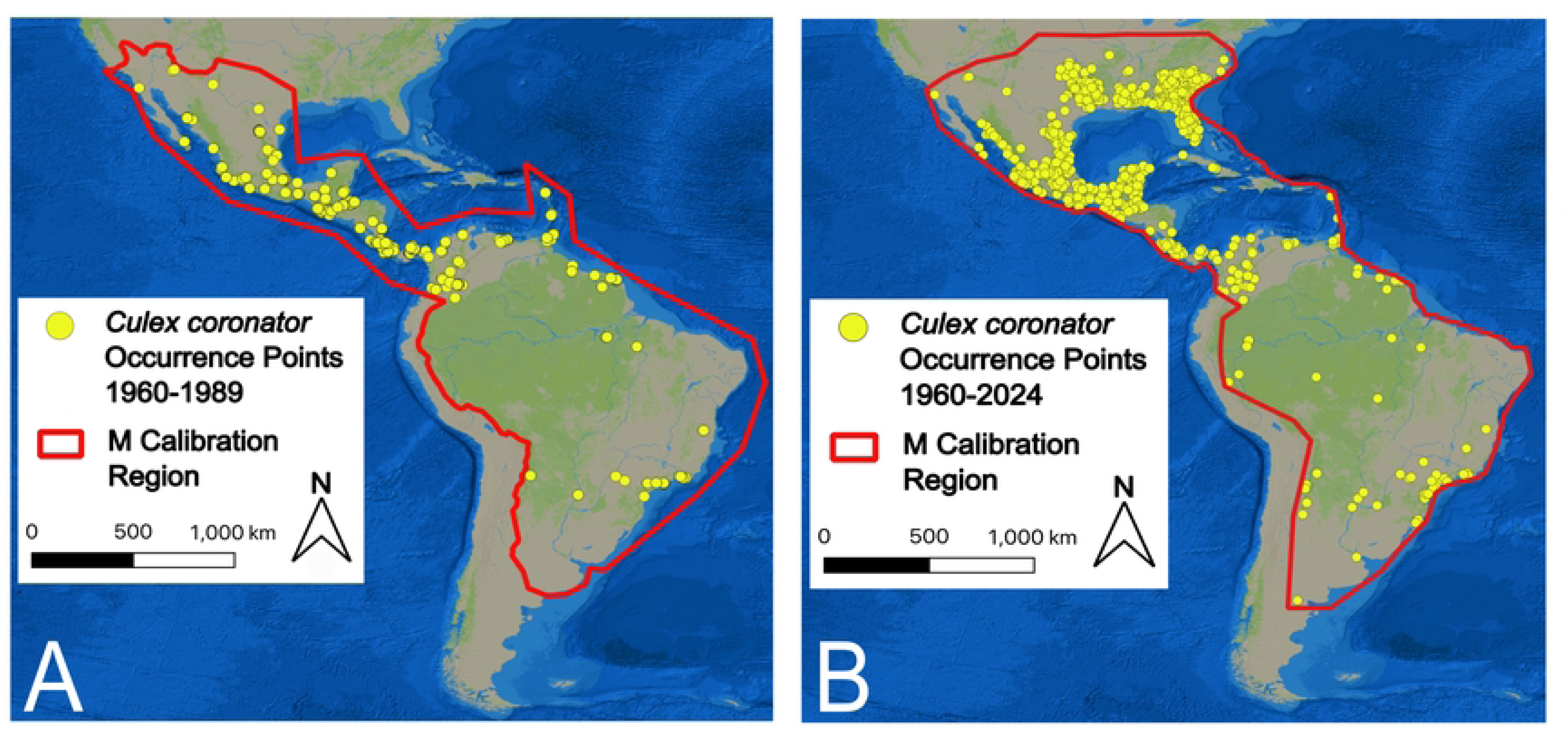
(A) Distribution of *Cx. coronator* georeferenced occurrence points from 1960 to 1989 used in historic model calibration and evaluation (yellow points) (B) Distribution of *Cx. coronator* georeferenced occurrence points from 1960-2024 used in contemporary model calibration and evaluation (yellow points). In both A & B, the red polygon depicts the **M**-calibration region. Base maps provided by the Environmental Systems Research Institute.

### Environmental Data

Bioclimatic variables at a 2.5-minute spatial resolution (∼5 km) were derived from downscaled Climate Research Unit (CRU TS.4.09) data, bias-corrected to WorldClim v2.1 data, available through the WorldClim website [67,68]. These variables are derived from combinations of monthly temperature and precipitation data and were calculated using the ‘dismo’ (v1.3-16) package [69,70]. Two bioclimatic datasets were developed for model calibration and prediction using monthly minimum and maximum temperature (tmin, tmax) and cumulative precipitation (prec) data from 1960 – 2024. Mean monthly climate variables were calculated for 1960 – 1989 and 2000 – 2024 using functions in the ‘raster’ package (v3.6-32) in R [70], and then used to generate bioclimatic variables for each time period. The same workflow was applied to the full 1960 – 2024 dataset to generate variables for the contemporary model.

For model calibration, bioclimatic variables from 1960–1989 were masked to the historical **M** calibration region (Figure 1A), while variables from 1960–2024 were masked to the contemporary **M**-region (Figure 1B). Next, multicollinearity among variables was assessed using variance inflation factor (VIF) values calculated with the *usdm* package in R [71]. Variables with VIF > 5 were not included within the same candidate model set. Based on this threshold, three candidate variable sets were generated for both the historical and contemporary model calibrations (Supplementary Tables 1&2).

To project the contemporary model to future climate conditions, we obtained bioclimatic variables derived from CMIP6 data for future climate projections for 2041 to 2060, downloaded through WorldClim website. These bioclimatic variables were derived from future climate projections generated by three models: Model for Interdisciplinary Research on Climate (MIROC) [72], NASA Goddard Institute for Space Studies (GISS) [73], and the Centro Euro-Mediterraneo sui Cambiamenti Climatic (CMCC) [74]. All projections were generated for the Shared Socioeconomic Projection 3 (SSP3) scenario [75], which represents a mid-range emission scenario characterized by regionally-focused climate policies and limited mitigation efforts.

### Model Calibration

Models were calibrated in the R package ‘ENMeval’ (v2.0.5.2) [76] using the ‘maxnet’ algorithm. Candidate models were generated for each of the three sets of environmental variables and included an internal 50% random subset of the occurrence data for training and testing. Models were run using a combination of feature classes (ie., linear [l], quadratic [q], product [p], linear+quadratic [lq], linear+product [lp], quadratic+product [qp], linear+quadratic+product [lqp]), no extrapolation, and 30,000 background points. Regularization multipliers ranged from 0.1 to 5, at intervals of 0.1 for the historic model and 0.25 for the contemporary model.

### Model Evaluation and Projection

We used an information criterion approach for model evaluation, ranking models from each candidate set from lowest to highest Akaike’s information criterion (AIC_c_) scores corrected for small sample sizes [77]. We also observed training and testing area under the curve (AUC) values, as well as the AUC difference to gauge model generalizability [77]. The best performing model was then run in Maxent (Version 3.4.1) [78] using the same parameters, but with 20 bootstrap replicates with 500 iterations each to derive standard deviation values and to observe variable contributions to model performance.

The historical model was used to project environmental suitability across the broader Gulf Coast study area under both historical (1960–1989) and recent (2000–2024) environmental conditions. To quantify changes in predicted suitability between these periods, the historical projection was subtracted from the recent projection. Finally, the contemporary model, calibrated using all available occurrence data from 1960 to 2024, was projected across the same study area and projected to future climate conditions.

## Results

Occurrence data for historical model calibration initially included 2,894 georeferenced records. After removing duplicates, 2,890 records remained, which were further reduced to 120 points following spatial thinning for model calibration. Model selection based on AIC_c_ rankings identified candidate set 1 as the best-performing variable set. The best performing model used linear, quadratic, and product ([lqp]) feature classes with a regularization multiplier of 0.6 (Table 1). This model had the lowest AIC_c_ score, with a corresponding AIC_c_ weight of 0.96, a training AUC of 0.805, and an AUC difference of 0.03, indicating good potential for generalization to other areas. A comparable model implemented in Maxent produced a similar mean AUC of 0.797 across 20 bootstrap replicates (Table 1). Standard deviation values across replicates were low, suggesting consistent predictions among model runs (Supplementary Figure 1a). Among the 9 bioclimatic variables included, mean diurnal range, precipitation of the coldest quarter, and precipitation seasonality contributed the greatest to model performance (Table 1; Figure 2).

**Figure 2.**
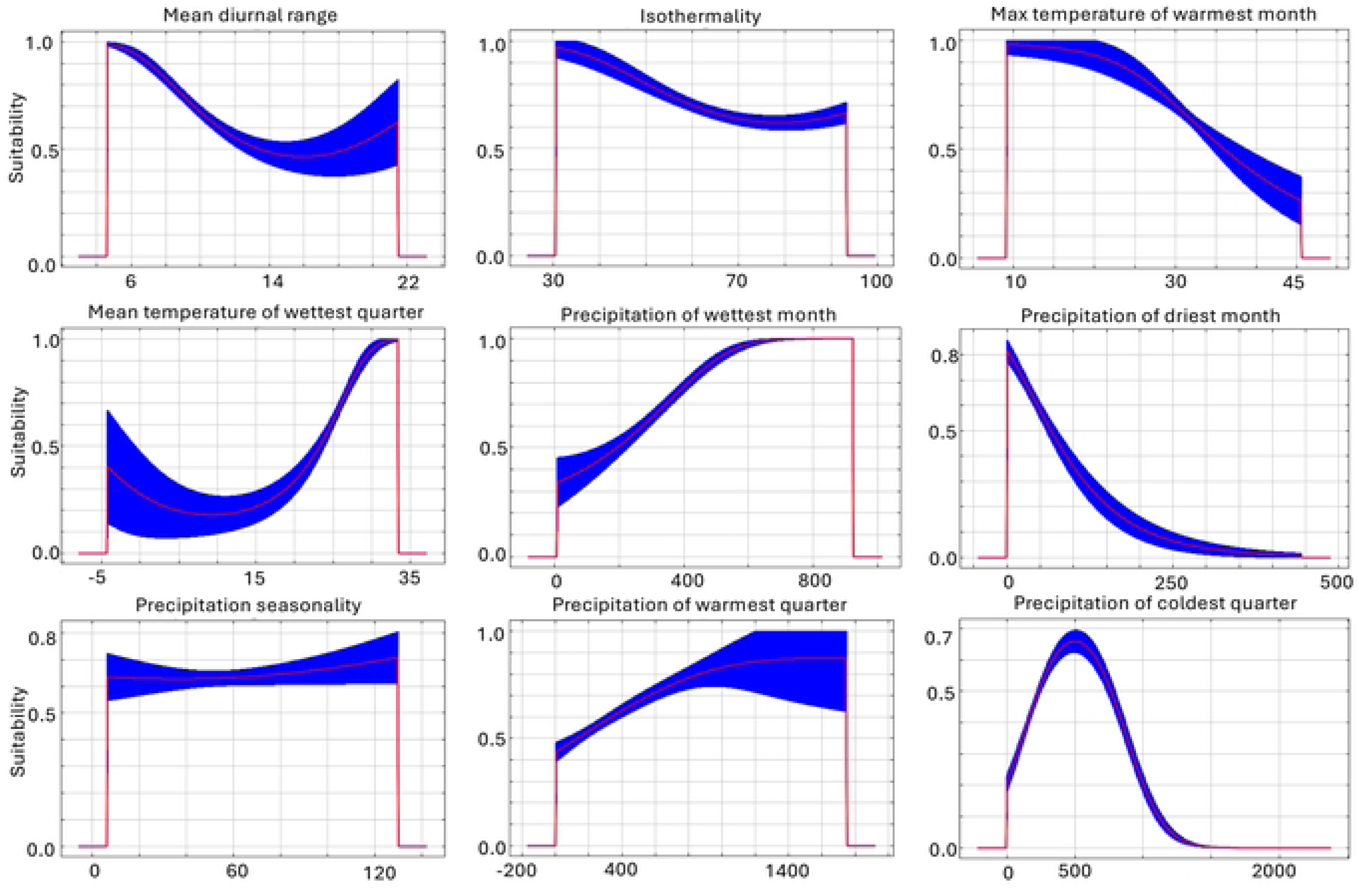
Response curves of bioclimatic variables used in the historical model. Temperature values are in units of degrees Celsius; precipitation is calculated in mm.

**Table 1.**
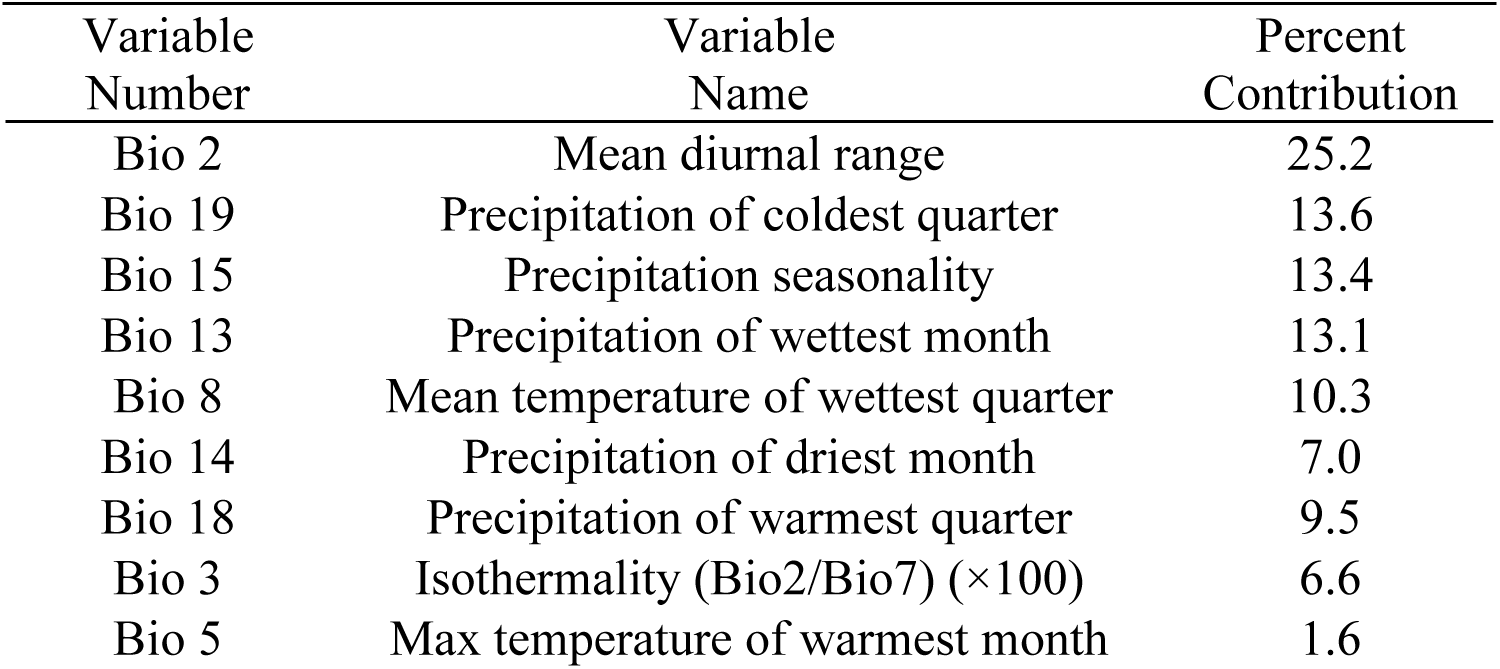
Bioclimatic variables included in the best-performing historic model and percent contribution to model performance.

| Variable<br>Number | Variable<br>Name | Percent<br>Contribution |
| --- | --- | --- |
| Bio 2 | Mean diurnal range | 25.2 |
| Bio 19 | Precipitation of coldest quarter | 13.6 |
| Bio 15 | Precipitation seasonality | 13.4 |
| Bio 13 | Precipitation of wettest month | 13.1 |
| Bio 8 | Mean temperature of wettest quarter | 10.3 |
| Bio 14 | Precipitation of driest month | 7.0 |
| Bio 18 | Precipitation of warmest quarter | 9.5 |
| Bio 3 | Isothermality (Bio2/Bio7) ($\times 100$) | 6.6 |
| Bio 5 | Max temperature of warmest month | 1.6 |

Marginal response curves showed non-linear responses between predicted suitability and the bioclimatic variables. Suitability was highest at lower values of mean diurnal range, or low daily temperature fluctuations, as well as moderate levels of precipitation during the coldest quarter. Response curves also showed a slight positive linear response to increasing precipitation seasonality (Figure 2).

Model projections for the historical (1960–1989; Figure 3A) and recent (2000–2024; Figure 3B) periods were broadly similar across parts of South and Central America and Mexico. However, increases in predicted suitability were observed along the eastern slopes of the Andes, as well as in French Guiana and northern Brazil. Across the U.S. Gulf Coast region, changes in predicted suitability between the two periods were generally low to moderate. The largest increases were concentrated in northern Mexico and southern Texas, followed by southern Louisiana, Alabama, Florida, and Cuba (Figure 4).

**Figure 3.**
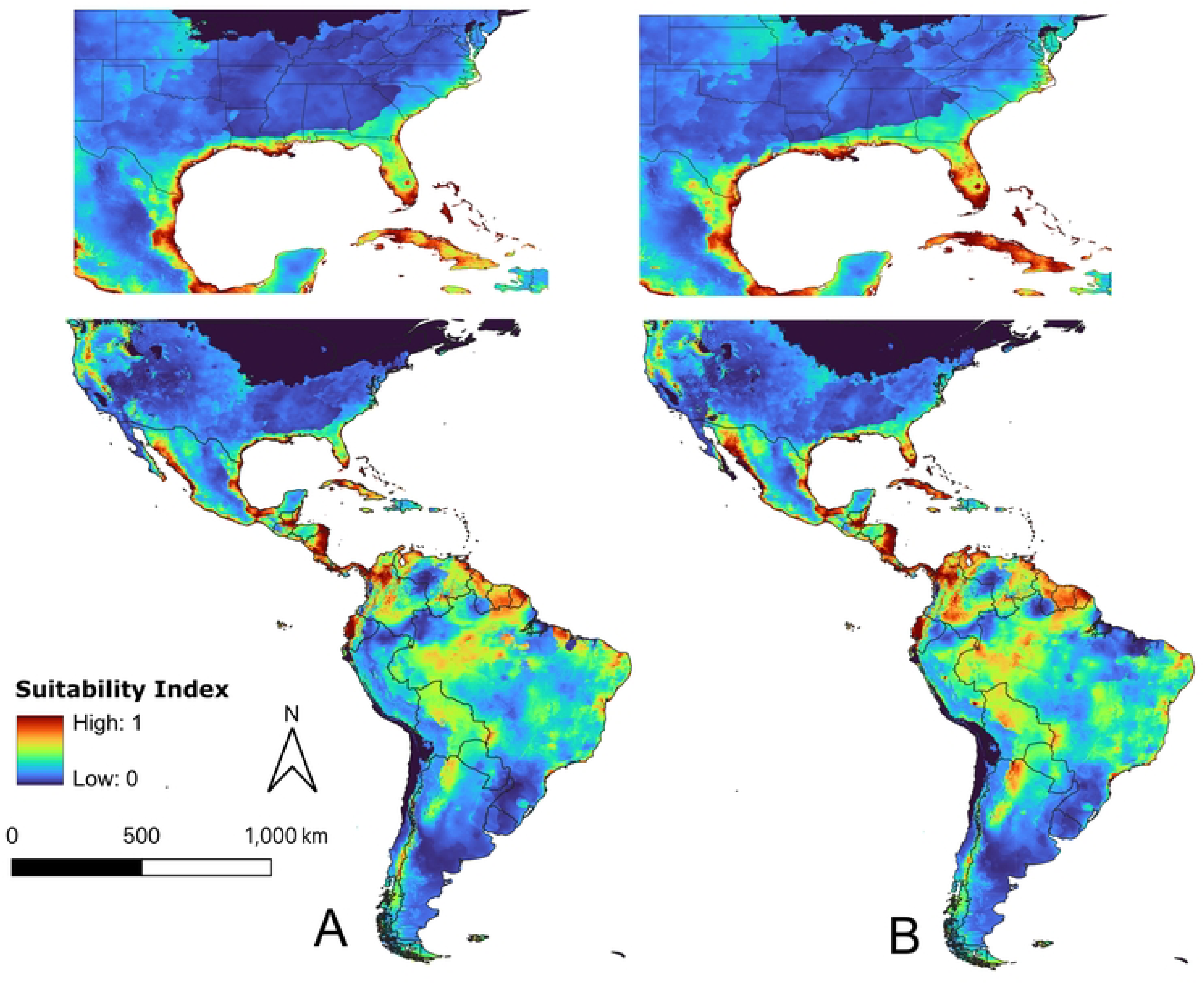
(A) Model predictions of past environmental suitability of *Cx. coronator*, 1960 to 1989, and (B) present environmental suitability 2000 to 2024 of *Cx. coronator*, calibrated using the occurrence data from 1960 to 1989.

**Figure 4.**
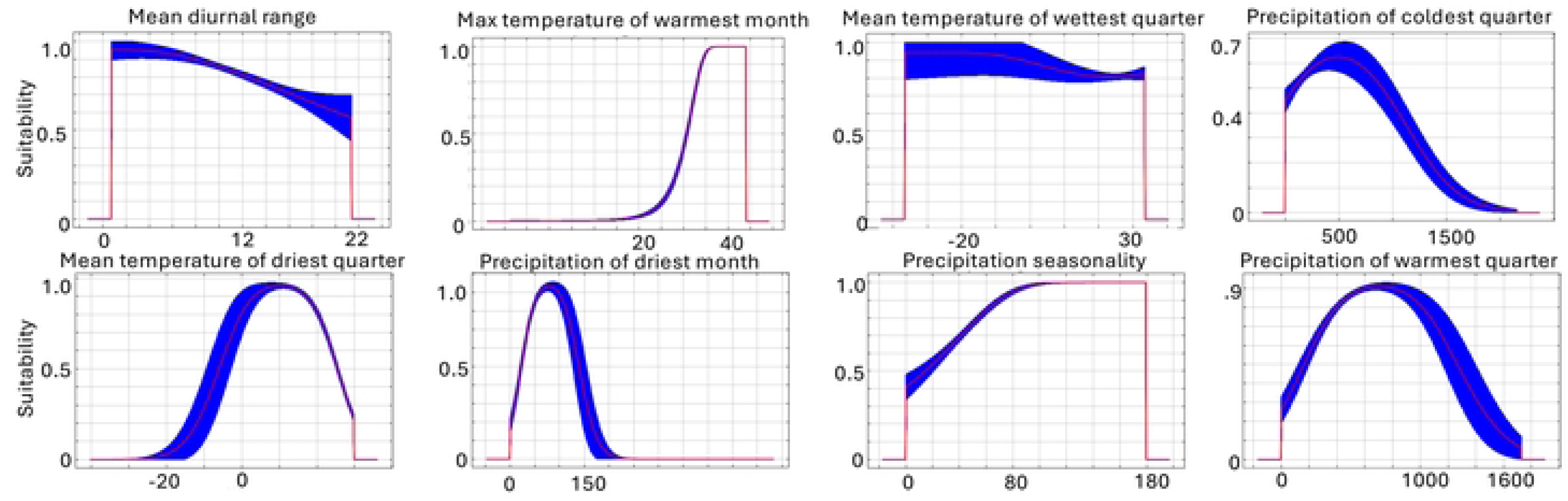
Change in predicted environmental suitability for *Cx. coronator* between historic (1960 to 1989) and recent (2000 to 2024) time periods. Areas with the greatest increases in predicted suitability are shown in lighter blue (difference = 0.99), areas with no changes are white (difference = 0), and areas with predicted the greatest decreased in suitability are brown (difference = −0.99). Yellow points represent the first occurrences of *Cx. coronator* in Louisiana and Mississippi recorded in 2004 and 2005, with changes in predicted suitability ranging from 0 to 0.37 at these locations.

At locations where *Cx. coronator* was first collected in 2004–2005, changes in predicted suitability ranged from 0 to 0.37 (Figure 4). A distinct band of increased suitability was also observed along the southern portion of the “Fall Line,” a geologic transition zone marked by increasing elevation that extends from Louisiana to South Carolina. In contrast, suitability at the historical range limit of *Cx. coronator* was predicted to be lower under recent climate conditions than under historical conditions, creating a gap across southern Texas. Despite this, the Texas coast remained highly suitable under recent climate conditions (Figure 3B).

Notably, projections to recent environmental conditions (2000–2024) predicted that environmentally suitable areas remained largely confined to the Gulf Coast, even though *Cx. coronator* has expanded its range widely across the humid subtropical region, suggesting that the model calibrated with historical data did not capture the full range of suitable environments for this species. Instead, many of these newly occupied areas were likely suitable in the past but were not accessible to the *Cx. coronator* at that time.

Occurrence data spanning the full study period (1960–2024) included 66,612 georeferenced *Cx. coronator* records. After removing duplicates, 59,870 records remained, which were reduced to 591 points following spatial thinning for model calibration. Model evaluation identified candidate set 1 as the best-performing variable set, with a [lqp] feature class and a regularization multiplier of 0.1. This model had the lowest AIC_c_ score, an AIC_c_ weight of 1, a training AUC of 0.821, and a low AUC difference of 0.025, indicating good generalization. The same model implemented in Maxent yielded a higher mean AUC of 0.906 across 20 bootstrap replicates, with low variation (maximum standard deviation = 0.174), suggesting consistent predictions. Eight bioclimatic variables were included, with mean temperature of the driest quarter, the maximum temperature of the warmest month, and precipitation of the driest month contributing most to model performance (Table 2).

**Table 2.** Bioclimatic variables included in the final contemporary model and average percent contribution to model performance.

| Variable Number | Variable Name | Percent Contribution |
| --- | --- | --- |
| Bio 9 | Mean temperature of driest quarter | 43.4 |
| Bio 5 | Max temperature of warmest month | 16 |
| Bio 14 | Precipitation of driest month | 11.3 |
| Bio 2 | Mean diurnal range | 11 |
| Bio 18 | Precipitation of warmest quarter | 8.6 |
| Bio 19 | Precipitation of coldest quarter | 3.8 |
| Bio 15 | Precipitation seasonality | 3.2 |
| Bio 8 | Mean temperature of wettest quarter | 2.6 |

Marginal response curves for the contemporary model also showed non-linear responses between suitability and key bioclimatic variables. Suitability was highest at low to moderate values of mean temperature during the driest quarter, declining at temperatures above ∼10°C (Figure 5). For the maximum temperature of the warmest month, predicted suitability increased from ∼20°C to ∼35°C, then dropped sharply at the upper limit of conditions represented in the calibration area. Similarly, suitability increased with precipitation of the driest month up to an apparent optimum (∼75 mm), after which it declined rapidly at higher values.

**Figure 5.**
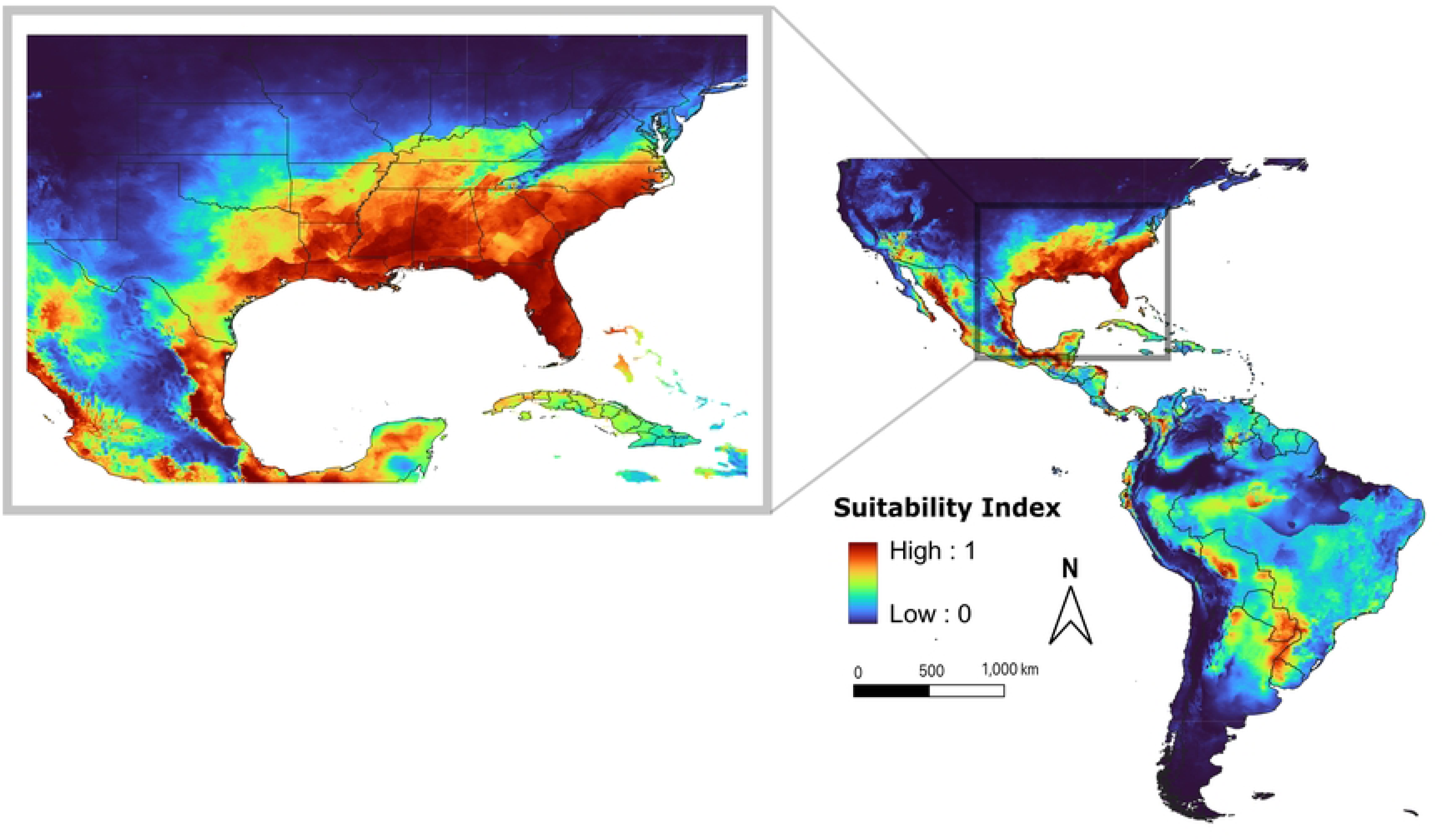
Response curves of bioclimatic variables used in the contemporary model. Temperature values are in units of degrees Celsius, and precipitation is measured in mm.

In contrast to the historical model, contemporary projections (1960–2024) predicted high environmental suitability across much of the humid subtropical region of the southern and eastern United States. These areas included the Gulf Coast and inland portions of Texas, Louisiana, Mississippi, Alabama, Georgia, and all of Florida (Figure 6). High suitability also extended northward along the Mid-Atlantic, including North and South Carolina, Virginia, and Maryland, before declining across Delaware and coastal New Jersey. Additional areas of high suitability were predicted inland across parts of Kentucky, except at higher elevations in the Appalachian Mountains. To the west, high suitability extended across Arkansas, Tennessee, Texas, and Oklahoma, with lower suitability predicted in Kansas and Missouri. Areas transitioning from humid subtropical to more arid environments were predicted to have low suitability for *Cx. coronator*. Low standard deviation values indicated strong agreement among model predictions (Supplementary Figure 1b).

**Figure 6.**
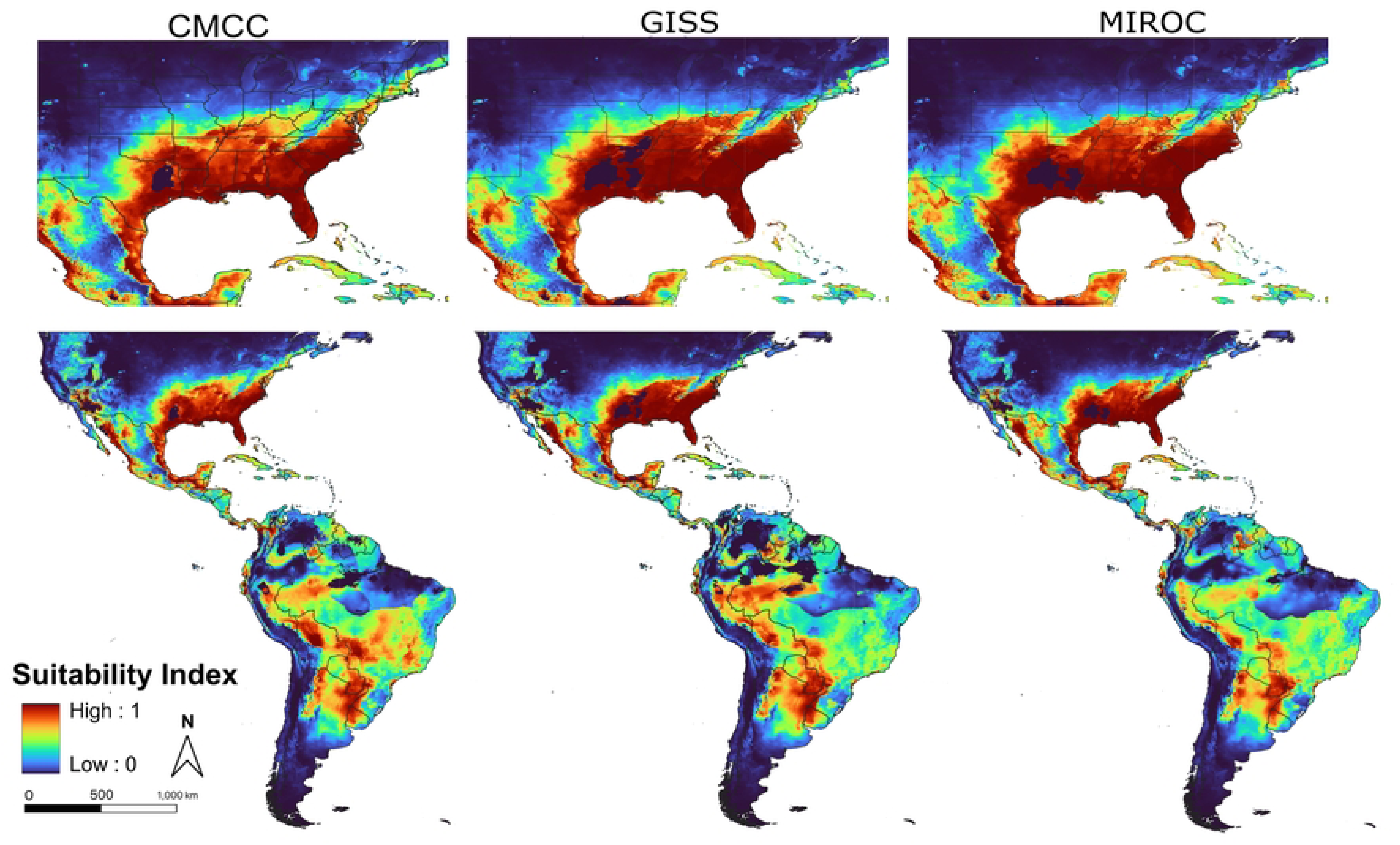
Contemporary predicted suitability for *Cx. coronator* across the full study period (1960 to 2024). The scale is from 0-1 representing relative suitability, and areas predicted to be highly suitable are in red, while areas with low predicted suitability are shown in blue.

Future projections for 2041–2060 under the SSP3 scenario were consistent across the three global climate models (CMCC, GISS, and MIROC), except for differences in areas of extrapolation (shown in black) where environmental conditions exceeded the calibration values and confidence was low (Figure 7). As expected, future model projections predicted a northward increase in suitability, primarily in the northeastern U.S., and highly suitable areas were more contiguous across the eastern U.S. Both contemporary and future projections also predicted high suitability in parts of California, although the species has not yet been reported west of Arizona (23).

**Figure 7.**
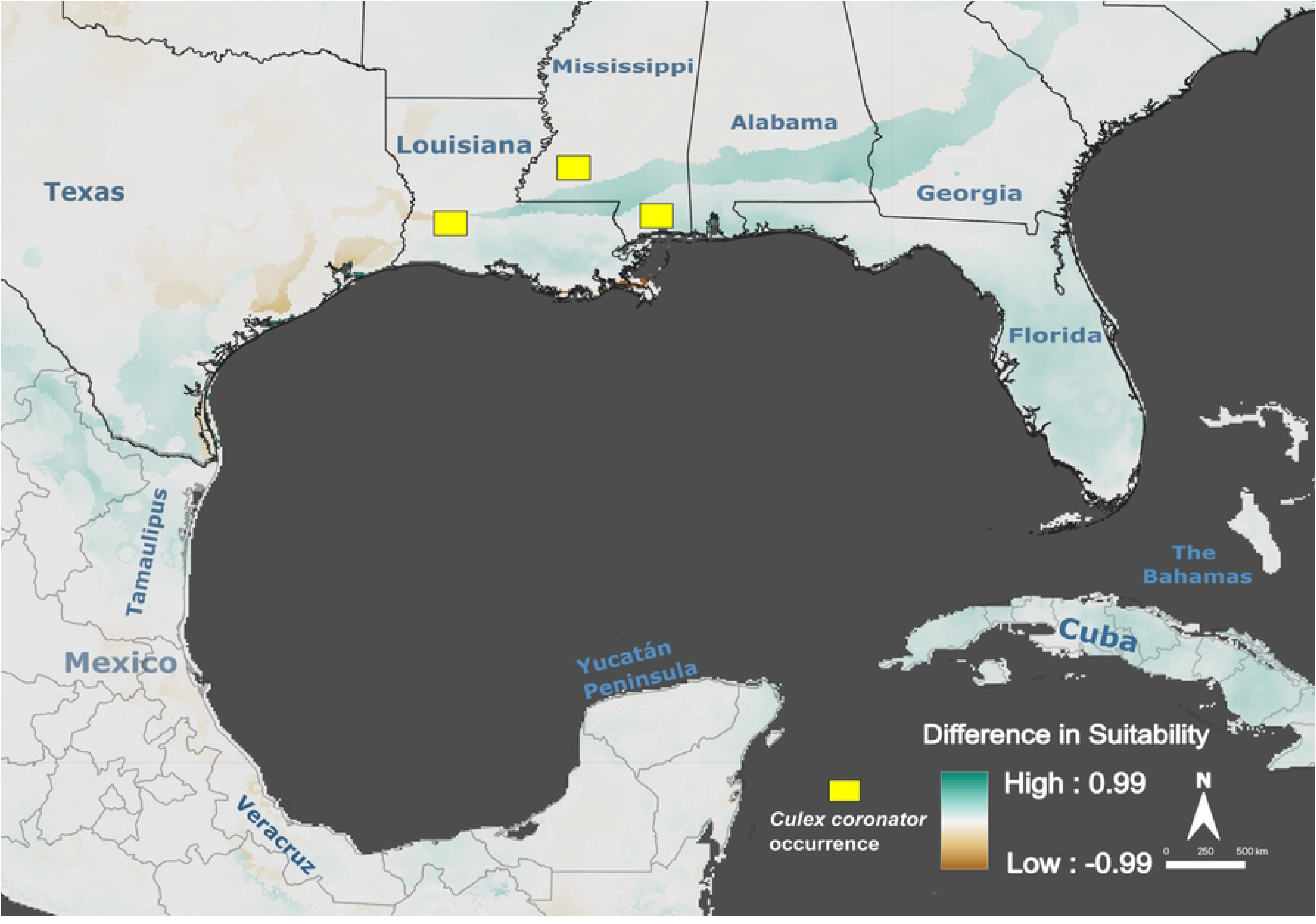
Future model predictions based on CMIP6 SSP3 scenario; A) Centro Euro-Mediterraneo sui Cambiamenti Climatici (CMCC)(79) B) NASA Goddard Institute for Space Studies (GISS)(78) C) Model for Interdisciplinary Research on Climate (MIROC)(77). Areas in red are predicted highly suitable; areas in blue have low predicted suitability; areas in black exceed combinations of environmental values available in model calibration, resulting in low confidence in predicted values.

## Discussion

This study modeled historical, contemporary, and future environmental suitability for *Cx. coronator* to evaluate patterns associated with its recent range expansion and to assess potential distributions under future climate conditions. Comparisons between projections for historical (1960–1989) and recent (2000–2024) conditions, based on models calibrated to the historical range, showed only moderate changes in suitability along the Gulf Coast, where *Cx. coronator* was first detected prior to broadscale range expansion in 2004. However, these models substantially underestimated the range of environments currently occupied by *Cx. coronator* across the humid subtropical eastern United States. In contrast, the contemporary model calibrated using the full occurrence dataset captured these environmental conditions and highlighted moisture availability as a potential key abiotic constraint. Together, these results underscore the value of updated prediction models to better anticipate the potential distribution of invasive mosquito species and inform mosquito surveillance and control efforts.

As expected, the contemporary model predicted widespread suitability across much of the humid subtropical eastern United States, with lower suitability at higher elevations in the Appalachian Mountains and at more northern latitudes with colder temperatures. Although the historical model included occurrences from humid subtropical regions of South America, the calibration relied heavily on tropical Central American records, which may have limited representation of cooler climates and contributed to underestimation of suitability in these regions. While thermal thresholds for *Cx. coronator* has not been reported, predicted temperature responses in the contemporary model align with values reported for other *Culex* species, particularly for the maximum temperature of the warmest month, where suitability increased between ∼20 and 35 °C before declining [79,80]. The ability of *Cx. coronator* to overwinter likely facilitates continued northward expansion. To date, the furthest northward confirmed collection of *Cx. coronator* is in Suffolk, Virginia [25]. However, model projections under future climate conditions show additional regions northward along the entirety of the Atlantic Coast and at higher elevations in the Appalachian Mountains that may be suitable for this species in the future.

Models consistently predicted low suitability in arid and semi-arid regions, suggesting that low moisture availability, rather than temperature alone, may represent a key abiotic constraint on *Cx. coronator* distributions. Here, the mean temperature of the driest quarter contributed most to model performance (43%), with suitability predicted highest near ∼20 °C and declining at warmer values, potentially reflecting increased desiccation opportunities for adults as well as eggs. While multiple *Culex* species are present in the western U.S., *Culex* eggs are particularly vulnerable to desiccation compared to some other mosquito genera [81]; for example, Vargas et al. (2014) found that *Cx. quinquefasciatus* eggs survive only a few hours under dry conditions. This sensitivity, combined with unmeasured biotic constraints, may limit westward expansion into hot, semi-arid environments, consistent with the lack of observed dispersal beyond *Cx. coronator’s* historic range.

Hot semi-arid climates also occur along the Gulf Coast between Texas and Louisiana further corresponding to the historical range limit of *Cx. coronator*. The contemporary model predicted low suitability in this region, suggesting that these conditions may have previously acted as a barrier to expansion. Although beyond the scope of inference from the models developed here, Connelly et al. (2016) hypothesized that extreme weather events, including hurricanes combined with overall warmer winter temperatures, may have facilitated movement of *Cx. coronator* across this corridor by creating temporary sunlit larval habitats supporting survival and reproduction [27]. The role of extreme weather events in range expansion has been highlighted in multiple systems [82,83] and warrants further investigation, given the number of mosquito species moving from historical distributions across the Gulf Coast region.

A key limitation of this study is continued potential for sampling bias of *Cx. coronator* occurrence data. However, the large number of georeferenced occurrence points available across broad areas of recent range expansion help to mitigate this problem. Further, SDMs do not account for biotic interactions with other species, including competition or predation [84], and they assume equilibrium with environmental conditions [85], which can be important components when considering successful movement and establishment of a species across a new area [86]. Despite these limitations, the use of SDMs to predict potential distributions under past, current and future climate conditions provided a useful tool to observe patterns across a time period of rapid range expansion for *Cx. coronator*.

Overall, this study utilized species distribution models to evaluate historic, current, and future abiotic conditions favorable for *Cx. coronator* across its known historical distribution and expanded range. More broadly, predicting potential distributions of invasive mosquito species enables operational control programs to prepare for potential establishment and to train personnel on taxonomic identification of these invasive species. These strategies will allow for a more robust, data-driven integrated mosquito management program that can target mosquito control efforts and continue to protect public health.

## Data Availability

All occurrence data and code used in the modeling and analyses are openly available without restriction in a public GitHub repository (https://github.com/Campbell-Lab-FMEL/Culex-coronator). Environmental data, occurrence data, and code are also archived without restriction in a Zenodo repository (https://doi.org/10.5281/zenodo.21169786). GBIF-mediated occurrence data were downloaded on 17 December 2024 and are available via GBIF at https://doi.org/10.15468/dl.wf953x.

## Acknowledgements

The authors would like to acknowledge mosquito control & public health personnel, entomologists, and community scientists who gather and publish the occurrence records. We also would like to extend our thanks to the World Climate Research Program, which, through its Working Group on Coupled Modeling, coordinated and promoted CMIP6.

---

Supplementary Figure 1a. Standard deviation of predicted suitability for historic model across 20 bootstrap replicates; light green indicates higher standard deviation values, and dark blue areas indicate areas with low standard deviation values.

Supplementary Figure 1b. Standard deviation of predicted suitability across 20 bootstrap replicates; light green indicates higher standard deviation values, and dark blue areas indicate areas with low standard deviation values.

